# FER-LIKE FE DEFICIENCY-INDUCED TRANSCRIPTION FACTOR (OsFIT) interacts with OsIRO2 to regulate iron homeostasis

**DOI:** 10.1101/2020.03.06.981126

**Authors:** Gang Liang, Huimin Zhang, Yang Li, Mengna Pu, Yujie Yang, Chenyang Li, Chengkai Lu, Peng Xu, Diqiu Yu

## Abstract

There are two Fe-uptake strategies for maintaining Fe homeostasis in plants. As a special graminaceous plant, rice applies both strategies. However, it remains unclear how these two strategies are regulated in rice. IRON-RELATED BHLH TRANSCRIPTION FACTOR 2 (OsIRO2) is critical for regulating Fe uptake in rice. In this study, we identified an interacting partner of OsIRO2, *Oryza sativa* FER-LIKE FE DEFICIENCY-INDUCED TRANSCRIPTION FACTOR (OsFIT), which encodes a bHLH transcription factor. The OsIRO2 protein is localized in the cytoplasm and nucleus, but OsFIT facilitates the accumulation of OsIRO2 in the nucleus. Loss-of-function mutations to *OsFIT* result in decreased Fe accumulation, severe Fe-deficiency symptoms, and disrupted expression of Fe-uptake genes. In contrast, *OsFIT* overexpression promotes Fe accumulation and the expression of Fe-uptake genes. Genetic analyses indicated that *OsFIT* and *OsIRO2* function in the same genetic node. Further analysis suggested that OsFIT and OsIRO2 form a functional transcription activation complex to initiate the expression of Fe-uptake genes. Our findings provide a mechanism understanding of how rice maintains Fe homeostasis.

**One-sentence summary:** OsFIT interacts with and facilitates the accumulation of OsIRO2 in the nucleus where the OsFIT-OsIRO2 transcription complex initiates the transcription of Fe deficiency responsive genes.

## INTRODUCTION

Iron (Fe) is necessary for plant growth and development because it is involved in many physiological and biochemical reactions. Although Fe is the second most abundant metal element in the earth crust, Fe availability is extremely low in highly alkaline soils, especially calcareous soils (Mori, 1999). Fe deficiency can cause serious agricultural problems, such leaf chlorosis as well as diminished plant growth and crop yield. Therefore, maintaining Fe homeostasis is important for ensuring plant growth and development.

Plants have evolved two strategies for increasing the efficiency of Fe uptake from soil (Marschner et al., 1986). Graminaceous plants apply strategy II, which involves the synthesis and secretion of mugineic acid family phytosiderophores (MAs) that solubilize and chelate Fe(III) in the rhizosphere for the subsequent absorption of MA-Fe(III) via plasma membrane transporters. In contrast, non-graminaceous plants employ strategy I, which requires the acidification of the rhizosphere to promote Fe release, the reduction of Fe(III) to Fe(II) at the root surface, and the subsequent uptake of Fe(II). Recent studies suggest that Arabidopsis also secretes Fe-chelating compounds, such as coumarins (Rodriguez-Celma and Schmidt, 2013; Fourcroy et al., 2014; Schmid et al., 2014; Siwinska et al., 2018; Tsai et al., 2018).

The key genes involved in both Fe-uptake strategies have been characterized in Arabidopsis and rice. Regarding Arabidopsis strategy I, the rhizosphere acidification in response to Fe deficiency is mediated by the plasma membrane H^+^-ATPase 2 (AHA2) (Santi and Schmidt, 2009), and the subsequent Fe(III) reduction and Fe(II) transport are mediated by FERRIC REDUCTASE 2 (FRO2) (Robinson et al., 1999) and IRON TRANSPORTER 1 (IRT1) (Connolly et al., 2002; Varotto et al., 2002; Vert et al., 2002). In rice, MAs are synthesized via the conversion of methionine to 2′-deoxymugineic acid in four sequential steps mediated by S-adenosylmethionine synthetase (SAMS), nicotianamine synthase (NAS), nicotianamine aminotransferase (NAAT), and deoxymugineic acid synthase (DMAS) (Bashir et al., 2006; Mori, 1999; Shojima et al., 1990). Additionally, TRANSPORTER OF MAs 1 (OsTOM1) mediates the efflux of MAs (Nozoye et al., 2011) and YELLOW STRIP LIKE 15 (OsYSL15) mediates the influx of MA-Fe(III) (Inoue et al., 2009; Lee et al., 2009). In addition to strategy II based on MAs, rice also partially employs strategy I involving the direct uptake of Fe(II) by OsIRT1 (Ishimaru et al., 2006).

Under Fe-deficient conditions, plants detect changes in the internal Fe concentration, after which the Fe uptake system is activated. Considerable progress has been made regarding the characterization of the Fe-deficiency-responsive signaling pathway in plants (Gao et al., 2019; Wu and Ling, 2019; Schwarz and Bauer, 2020). In Arabidopsis, BTS is a putative Fe sensor because its hemerythrin motifs bind to Fe and it contains a Really Interesting New Gene (RING) domain associated with ubiquitination activity. Additionally, BTS negatively regulates Fe homeostasis by interacting with AtbHLH105 and AtbHLH115 to induce their degradation (Selote et al., 2015). In response to Fe deficiency, Arabidopsis bHLH IVc transcription factors (TFs) (AtbHLH34, AtbHLH104, AtbHLH105, and AtbHLH115) activate the expression of bHLH Ib genes (*AtbHLH38, AtbHLH39, AtbHLH100*, and *AtbHLH101*) and *AtPYE* (Zhang et al., 2015; Li et al., 2016; Liang et al., 2017). Moreover, bHLH Ib TFs interact with AtFIT to activate the expression of strategy I genes *AtIRT1* and *AtFRO2* (Yuan et al., 2008; Wang et al., 2013). A similar regulatory network also occurs in rice. As the homologs of BTS, OsHRZ1 and OsHRZ2 were identified as putative rice Fe sensors that also contain hemerythrin motifs and a RING domain (Kobayashi et al., 2013). Previous studies revealed that OsHRZ1 negatively regulates Fe homeostasis by mediating the degradation of OsPRI1, OsPRI2, and OsPRI3 (orthologs of Arabidopsis bHLH IVc), which positively regulate Fe homeostasis by directly targeting *OsIRO2* (homolog of Arabidopsis bHLH Ib) and *OsIRO3* (homolog of *AtPYE*) (Zhang et al., 2017, 2020; Kobayashi et al., 2019). Both *OsIRO2* and *OsIRO3* exhibit upregulated expression in response to Fe deficiency and their products positively and negatively regulate Fe homeostasis, respectively (Ogo et al., 2006, 2007; Zheng et al. 2010). As the rice ortholog of Arabidopsis bHLH Ib, OsIRO2 controls the expression of strategy II genes *OsNAS1, OsNAS2, OsNAAT1, OsDMAS1*, and *OsYSL15* (Ogo et al., 2007).

In this study, we functionally characterize OsFIT, which interacts with OsIRO2 and positively regulates rice Fe homeostasis. OsIRO2 recognizes and OsFIT activates the promoters of their target genes, such as *OsNAS2* and *OsYSL15*. Our data suggest that OsFIT and OsIRO2 function as a transcription complex to regulate Fe homeostasis.

## RESULTS

### OsFIT physically interacts with OsIRO2

Yeast two-hybrid (Y2H) assays were used to identify the interacting partners of OsIRO2. Because of the considerable self-activation of the full-length OsIRO2 (Supplemental Figure S1A), we used a truncated OsIRO2 (i.e., OsIRO2-N), in which 98 amino acids were deleted from the C terminus, as the bait for the Y2H screening of an iron-depleted rice cDNA library. Six of 112 positive clones contained the same prey protein, bHLH156 (Os04g0381700) (Supplemental Table S1), which was named *Oryza sativa* FER-LIKE FE DEFICIENCY-INDUCED TRANSCRIPTION FACTOR (OsFIT) because its protein sequence is similar to AtFIT (Supplemental Figure S1B). The full-length *OsFIT* coding region was cloned, after which the interaction between OsFIT and OsIRO2 was confirmed in yeast cells (Figure 1A).

**Figure 1.**
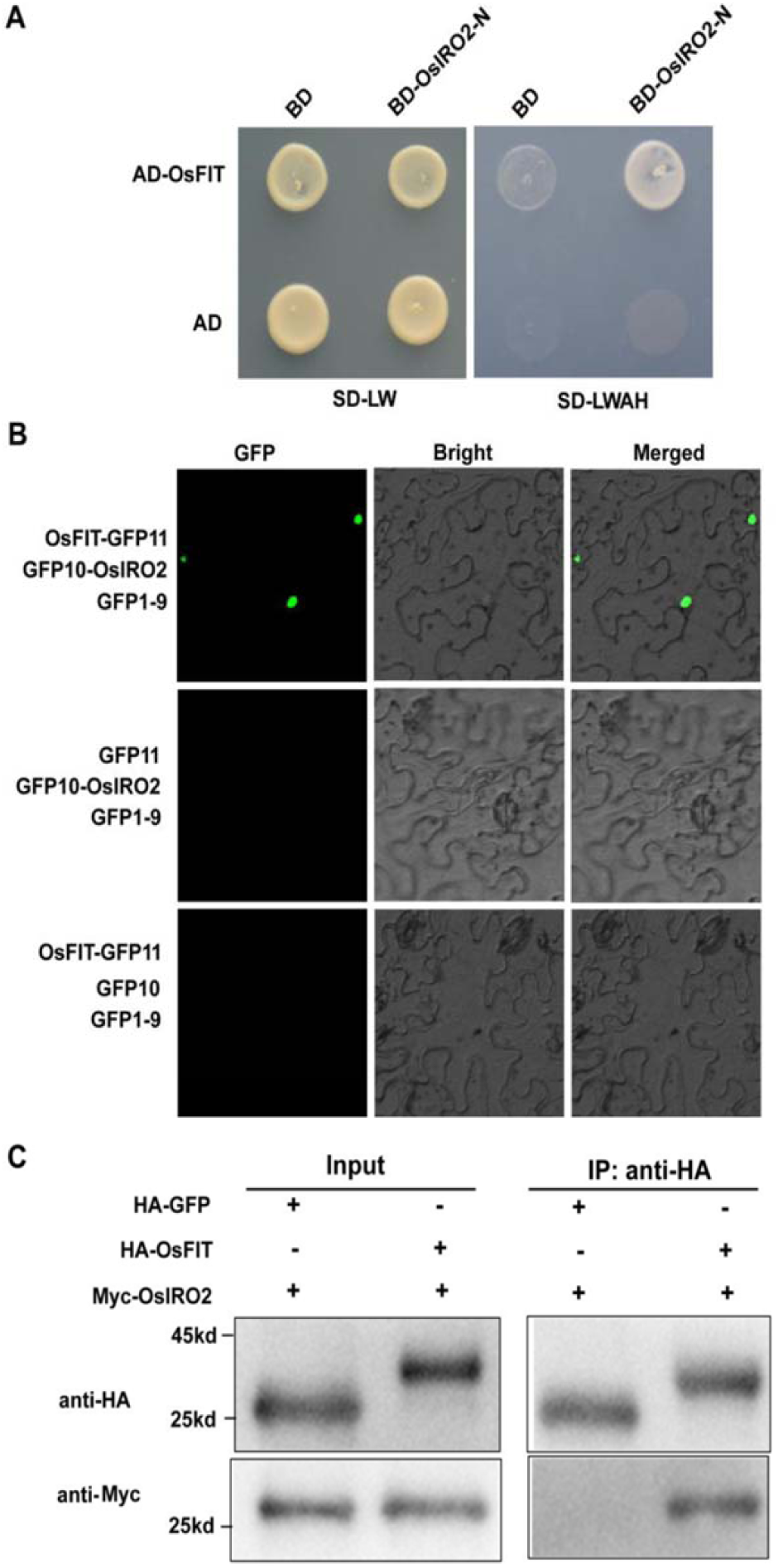
OsFIT physically interacts with OsIRO2. **(A)** Yeast two-hybrid analysis of the interaction between OsIRO2 and OsFIT. Yeast cotransformed with different BD and AD plasmid combinations was spotted on synthetic dropout medium lacking Leu/Trp (SD–W/L) or Trp/Leu/His/Ade (SD–W/L/H/A). **(B)** Fluorescence complementation between OsIRO2 and OsFIT. Three different combinations (GFP1-9/GFP10-OsIOR2/OsFIT-GFP11, GFP1-9/GFP10/OsFIT-GFP11, and GFP1-9/GFP10-OsIOR2/GFP11) were co-expressed respectively in tobacco leaves. **(C)** Co-IP analysis of the interaction between OsIRO2 and OsFIT. Total proteins from different combinations (HA-GFP/Myc-OsIRO2 and HA-OsFIT/Myc-OsIRO2) were immunoprecipitated with anti-Myc followed by immunoblotting with the indicated antibodies.

To further verify that OsIRO2 interacts with OsFIT in plant cells, we employed the tripartite split-GFP system (Liu et al., 2018). The full-length OsIRO2 protein was fused to the GFP10 fragment (GFP10-OsIRO2), whereas the full-length OsFIT was fused to the GFP11 fragment (OsFIT-GFP11). When GFP10-OsIRO2 and OsFIT-GFP11 were transiently co-expressed with GFP1-9 in tobacco leaves, the GFP signal was detected in the nucleus (Figure 1B). In contrast, a GFP signal was undetectable in cells containing control vectors.

We next conducted the co-immunoprecipitation assays to confirm the interaction between OsIRO2 and OsFIT *in planta*. The Myc-tagged OsIRO2 and the HA-tagged OsFIT were transiently co-expressed in tobacco leaves. The proteins were incubated with the anti-Myc antibody and A/G-agarose beads and then separated by SDS-PAGE for immunoblotting with the anti-HA and anti-MYC antibodies. Consistent with the results of the Y2H and tripartite split-GFP assays, OsIRO2 and OsFIT were detected in the same protein complex (Figure 1C). These data suggest that OsIRO2 physically interacts with OsFIT.

### OsFIT promotes the nuclear accumulation of OsIRO2

To investigate the effects of OsFIT on Fe homeostasis, we determined whether *OsFIT* expression is responsive to Fe deficiency. Wild-type seedlings grown in 0.1 mM Fe(III) solution (+Fe) for 5 days were shifted to a +Fe or Fe-free solution (−Fe) for 5 days. The roots and shoots were harvested separately for RNA extraction. A quantitative real-time polymerase chain reaction (qRT-PCR) assay was conducted to analyze the expression of both *OsIRO2* and *OsFIT* (Figure 2A). In agreement with previous studies (Ogo et al., 2006, 2007), *OsIRO2* expression increased in the roots and shoots under −Fe conditions. Similarly, *OsFIT* expression in the roots and shoots was also upregulated under −Fe conditions. Further analysis indicated that *OsFIT* is expressed preferentially in the roots rather than shoots (Figure 2B).

**Figure 2.**
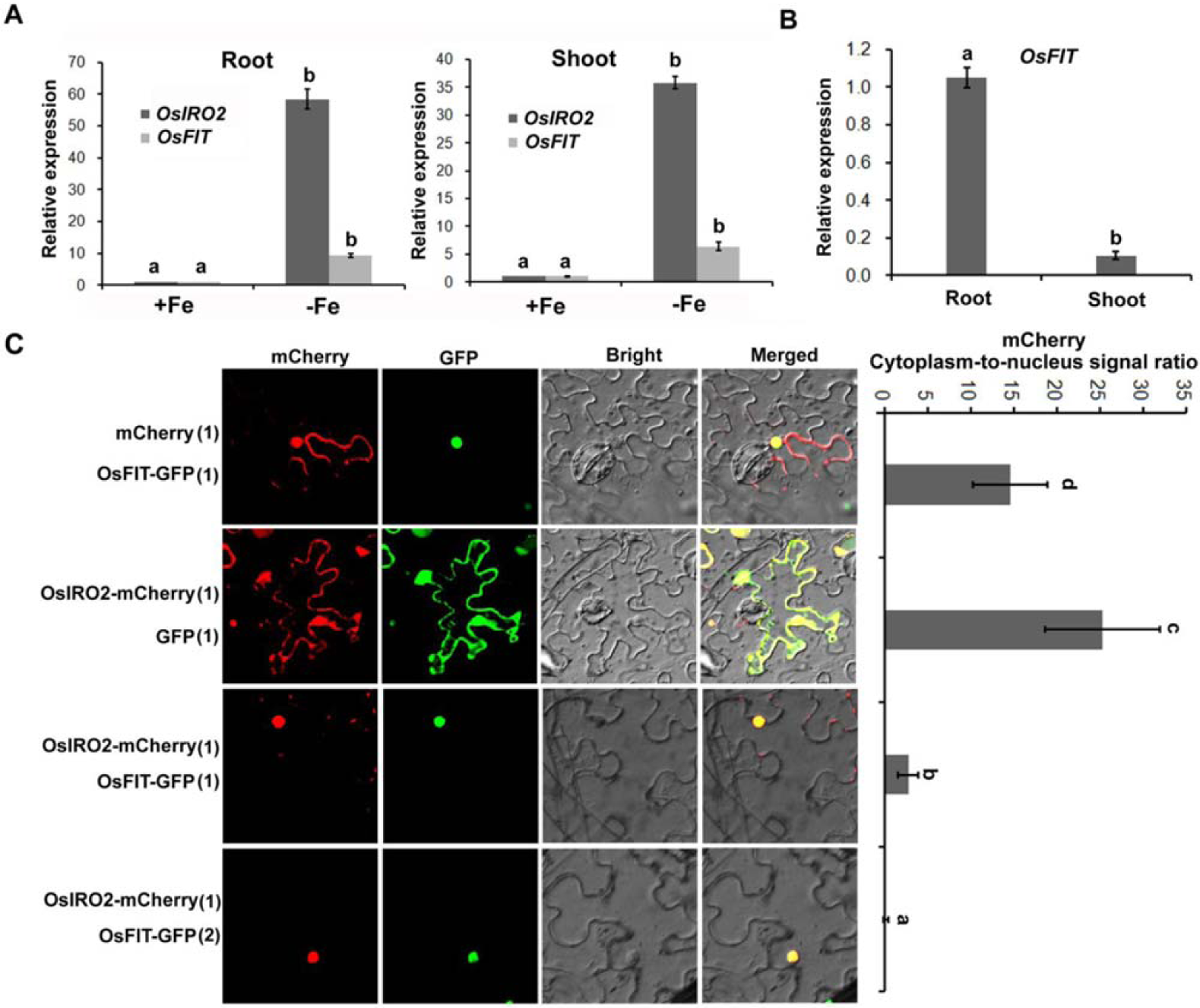
OsFIT facilitates the accumulation of OsIRO2 in the nucleus. **(A)** Response of *OsFIT* to Fe deficiency. Four-day-old seedlings germinated in wet paper were grown in 0.1 mM Fe (III) solution (+Fe) for 5 days and then transferred to +Fe or Fe free solution (–Fe) for 5 days. **(B)** *OsFIT* expression in the roots and shoots. Four-day-old seedlings germinated in wet paper were grown in 0.1 mM Fe (III) solution (+Fe) for 10 days. (A) and (B) Roots and shoots were harvested separately and used for RNA extraction and qRT-PCR. Data represent means ± SD (*n* = 3). **(C)** Subcellular localization. Different combinations of OsIRO2-mCherry, OsFIT-GFP, free GFP or free mCherry were expressed transiently in tobacco cells. The numbers in the parenthesis indicate the relative proportion of agrobacteria in each combination. Quantification of subcellular distribution of the mCherry tagged proteins are shown on the right. Data represent means ± SD (*n* = 10). (A-C) Different letters above each bar indicate statistically significant differences as determined by one-way ANOVA followed by Tukey’s multiple comparison test (P < 0.05).

To further examine the spatiotemporal *OsFIT* expression pattern, a construct comprising a β-glucuronidase (GUS) gene under the control of the 3.2-kb putative promoter upstream of *OsFIT* was prepared and transferred into wild-type rice. The GUS stain was detected in the roots and shoots and was more intense under -Fe conditions than under +Fe conditions (Supplemental Figure S2).

Subsequently, we examined the subcellular localization of OsFIT and OsIRO2 (Figure 2C). The full-length OsIRO2 was fused in frame with the mCherry and the full-length OsFIT with the GFP under the control of the CaMV 35S promoter. When transiently expressed in tobacco cells, OsFIT-GFP was exclusively targeted to the nucleus, whereas OsIRO2-mCherry was mainly localized in the cytoplasm. Their different subcellular localizations and interaction in the nucleus prompted us to investigate whether OsFIT influences the localization of OsIRO2. When equal amounts of OsIRO2-mCherry and OsFIT-GFP were mixed and transiently co-expressed, the fluorescence signal of cytoplasm OsIRO2-mCherry decreased significantly. As the proportion of OsFIT-GFP increased, almost all OsIRO2-mCherry concentrated in the nucleus. These results suggested that OsFIT promotes the nuclear accumulation of OsIRO2.

### Loss-of-function of *OsFIT* impairs tolerance to Fe limitation

To investigate the physiological functions of OsFIT, the CRISPR/Cas9 editing system was employed to edit the *OsFIT* gene. Two target sites within the bHLH domain were designed and respectively integrated into the CRISPR/Cas9 editing vector (Supplemental Figure S3A), which were introduced into wild-type rice via *Agrobacterium tumefaciens*-mediated transformation. Two homozygous mutants (*fit-1* and *fit-2*) were identified by a sequencing analysis (Figure 3A). Both mutants contain a T insertion in the bHLH domain, which introduced a STOP codon in the *fit-1* and caused the frame-shift mutation in the *fit-2* (Supplemental Figure S3B). Further expression analysis indicated that the *OsFIT* expression was lower in both mutants than in the wild-type (Supplemental Figure S3C).

**Figure 3.**
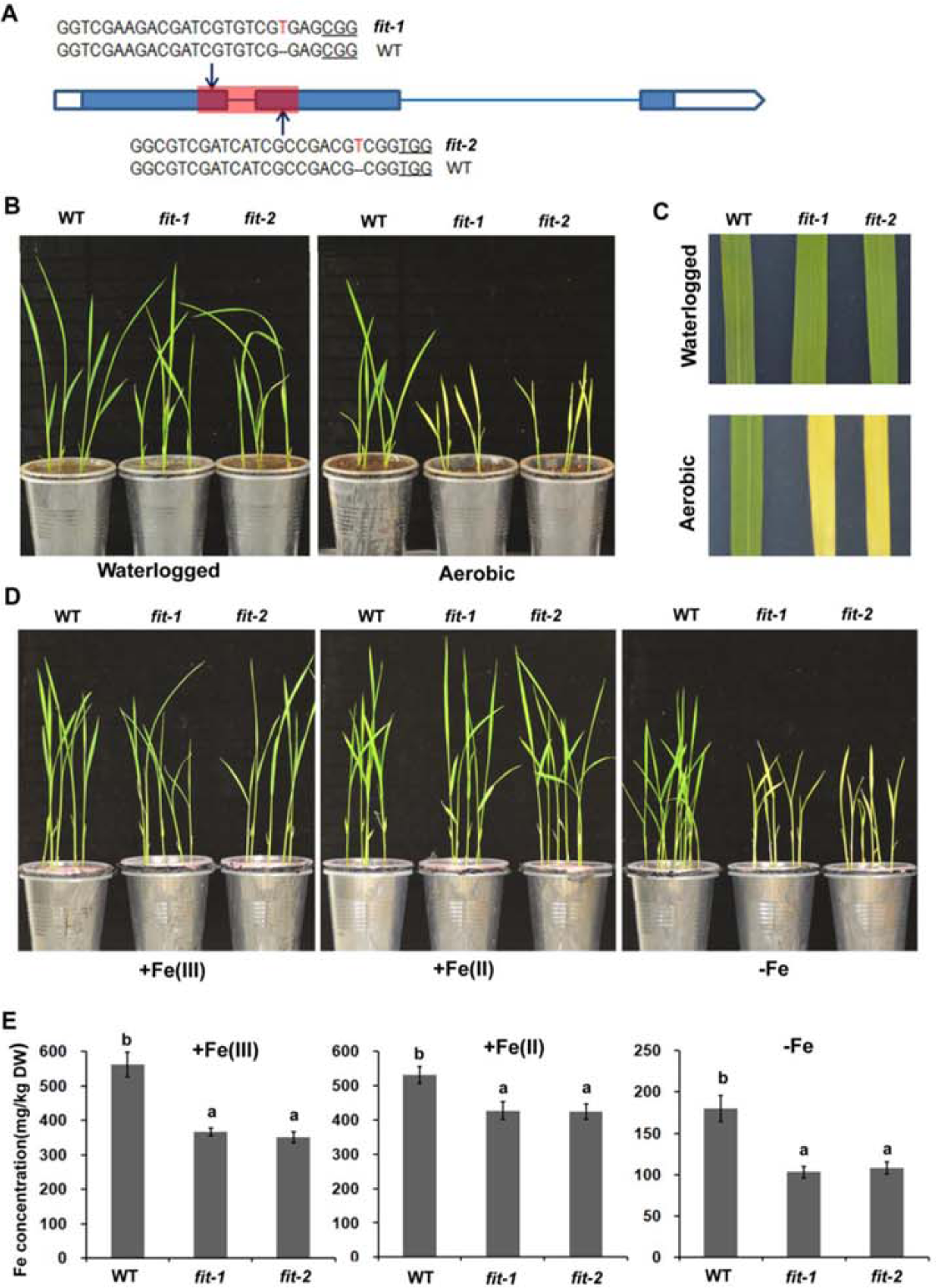
Phenotypes of *fit* mutants. **(A)** CRISPR/Cas9-edited *fit* mutants. The underlined three letters indicate the PAM region. Arrows indicate the positions of single guide RNAs. The red letter indicates the 1bp insertion. The red bar indicates the bHLH domain. **(B)** Growth of *fit* mutant seedlings under aerobic conditions and water logged. Two-week-old seedling are shown. **(C)** The third leaves of seedlings in (B). **(D)** Growth of *fit* mutant seedlings. Seeds were germinated on wet paper for four days. For Fe (III) and Fe (II) growth, four-day-old seedlings germinated on wet paper were shifted in 0.1 mM Fe(III) and Fe (II) solution respectively for 10 days. For –Fe growth, four-day-old seedlings germinated on wet paper were shifted to +Fe for 1 day and then transferred to –Fe for 9 days. **(E)** Fe concentration of shoots in (D). Data represent means ± SD (*n* = 3). Different letters above each bar indicate statistically significant differences as determined by one-way ANOVA followed by Tukey’s multiple comparison test (P < 0.05).

When the *fit* mutants were grown in aerobic soil, the plants exhibited delayed development and severe chlorosis (Figure 3B, C; Supplemental Figure S3D). When grown in waterlogged soil, no visible phenotypic differences were observed between the *fit* mutants and wild-type plants (Figure 3B, C). Rice mainly uses strategy II to obtain Fe in aerobic soil lacking soluble Fe(II) (Liu et al., 2019). We speculated that the strategy II components were damaged in the *fit* mutants. To confirm our hypothesis, we performed hydroponic experiments involving a nutrient solution with various Fe contents [+Fe(III), 0.1 mM EDTA-Fe(III); +Fe(II), 0.1 mM EDTA-Fe(II); and −Fe, Fe free]. Regardless of the presence of Fe(III) or Fe(II), the *fit* mutants and the wild-type plants developed similarly; however, under −Fe conditions, the *fit* mutants exhibited Fe-deficiency hypersensitive phenotypes (i.e., very poor growth and extensive leaf chlorosis) (Figure 3D). Furthermore, a comparison of Fe concentrations indicated that the *fit* mutants accumulated less Fe than the wild-type plants regardless of Fe status (Figure 3E).

### Loss-of-function mutations to *OsFIT* disrupt the expression of Fe homeostasis-associated genes

To evaluate the effect of *OsFIT* mutations on the rice Fe-uptake system, we performed an Affymetrix GeneChip analysis. Nine-day-old seedlings grown in +Fe solution were shifted to +Fe or −Fe solution for 5 days. The shoots and roots were harvested separately for a GeneChip analysis (Figure 4A; Supplemental Figure S4A). The results revealed that among 741 genes upregulated by Fe depletion in the wild-type roots, 367 (50%) were downregulated in the *fit-1* mutant relative to the wild-type expression level. Moreover, of the 845 genes downregulated by Fe depletion in the wild-type roots, 465 (55%) were upregulated in the *fit-1* mutant (Figure 4B). In contrast, the gene expression levels were less affected in the shoots (Supplemental Figure S4B). We noted that the root expression levels of a group of genes involved in Fe uptake were substantially lower in the *fit-1* mutant than in the wild-type control (Table 1). These genes included strategy I genes (*OsIRT1* and *OsHA1*) and strategy II genes (*OsYSL9*/*15*/*16, OsTOM1, OsENA1*, and *OsZIFL9*). The genes associated with NA and DMA syntheses (*OsMTN, OsAPT1, OsMTK1, OsIDI1/2/4, OsDEP, OsFDH, OsNAS1, OsNAS2*, and *OsDMAS1*) whose expression levels were upregulated by Fe deficiency were expressed at lower levels in the *fit-1* mutant compared with the wild-type control. Previous studies confirmed that OsNAAT1 is a key enzyme for DMA synthesis in rice (Cheng et al., 2007; Inoue et al., 2008). Because of a lack of *OsNAAT1* probe in the GeneChip, we analyzed *OsNAAT1* expression via a qRT-PCR assay (Supplemental Figure S4C). The expression of *OsNAAT1* was downregulated in the *fit* mutants. Moreover, several Fe-uptake genes, such as *OsIRT1, OsENA1, OsENA2, OsTOM1, OsYSL15*, and *OsYSL16*, which were induced by Fe deficiency, exhibited downregulated expression in the *fit-1* mutants relative to the wild-type expression levels. Additionally, among the genes induced by Fe deficiency, *OsMIR, OsIMA1, OsNRAMP1, OsIRT2, OsIRO2*, and *OsIRO3* were more highly expressed in the *fit-1* mutant than in the wild-type control. These data suggest that the loss-of-function mutations to *OsFIT* disrupt the expression of genes associated with Fe homeostasis.

**Table 1.** Representative Fe deficiency responsive genes affected by OsFIT.

| ID | Root |  |  |  | Shoot |  |  |  | WT | *fit-1* | WT | *fit-1* | Annotation |
| --- | --- | --- | --- | --- | --- | --- | --- | --- | --- | --- | --- | --- | --- |
|  | WT | WT | *fit-1* | *fit-1* | WT | WT | *fit-1* | *fit-1* | Root | Root | Shoot | Shoot |  |
|  | +Fe | -Fe | +Fe | -Fe | +Fe | -Fe | +Fe | -Fe | -Fe/+Fe | -Fe/+Fe | -Fe/+Fe | -Fe/+Fe |  |
| Strategy I |  |  |  |  |  |  |  |  |  |  |  |  |  |
| Os03g0667500 | 11.79 | 14.90 | 11.32 | 13.10 | 8.56 | 8.88 | 8.57 | 8.66 | 3.12 | 1.78 | 0.32 | 0.09 | *OsIRT1* |
| Os05g0319800 | 2.09 | 5.97 | 2.08 | 2.04 | 2.08 | 2.09 | 5.27 | 2.26 | 3.88 | -0.04 | 0.01 | -3.02 | *OsHA1* |
| Strategy II |  |  |  |  |  |  |  |  |  |  |  |  |  |
| Os04g0542200 | 10.19 | 12.05 | 11.05 | 10.52 | 10.66 | 9.99 | 10.81 | 9.74 | 1.85 | -0.53 | -0.67 | -1.07 | *OsYSL9* |
| Os02g0650300 | 10.71 | 16.07 | 4.72 | 7.58 | 4.38 | 7.60 | 3.39 | 1.87 | 5.36 | 2.86 | 3.22 | -1.53 | *OsYSL15* |
| Os04g0542800 | 12.49 | 14.38 | 13.71 | 13.89 | 13.17 | 13.01 | 12.83 | 12.47 | 1.89 | 0.18 | -0.16 | -0.36 | *OsYSL16* |
| Os11g0134900 | 8.24 | 12.69 | 1.87 | 2.26 | 1.88 | 3.86 | 1.90 | 1.89 | 4.46 | 0.39 | 1.98 | -0.01 | *OsTOM1* |
| Os11g0151500 | 6.80 | 10.34 | 4.49 | 4.58 | 2.12 | 4.51 | 2.10 | 2.37 | 3.54 | 0.09 | 2.39 | 0.27 | *OsENA1* |
| Os12g0132500 | 9.85 | 14.54 | 4.68 | 4.70 | 3.49 | 6.62 | 3.63 | 3.49 | 4.69 | 0.02 | 3.13 | -0.14 | *OsZIFL9* |
| DMA synthesis |  |  |  |  |  |  |  |  |  |  |  |  |  |
| Os06g0112200 | 11.72 | 13.40 | 11.73 | 11.69 | 11.23 | 11.32 | 11.14 | 11.10 | 1.68 | -0.04 | 0.09 | -0.04 | *OsMTN* |
| Os12g0589100 | 13.81 | 15.47 | 13.80 | 13.88 | 13.57 | 13.79 | 13.49 | 13.63 | 1.66 | 0.08 | 0.22 | 0.14 | *OsAPT1* |
| Os04g0669800 | 12.45 | 14.18 | 12.23 | 12.55 | 12.31 | 12.61 | 12.30 | 12.34 | 1.74 | 0.31 | 0.31 | 0.05 | *OsMTK1* |
| Os11g0216900 | 9.91 | 12.11 | 9.68 | 10.23 | 11.61 | 11.63 | 11.47 | 11.56 | 2.20 | 0.55 | 0.02 | 0.09 | *OsIDI2* |
| Os11g0484000 | 12.62 | 15.46 | 12.19 | 12.49 | 13.21 | 13.26 | 13.05 | 13.27 | 2.83 | 0.29 | 0.05 | 0.21 | *OsDEP* |
| Os03g0161800 | 12.41 | 13.88 | 12.30 | 12.26 | 12.95 | 12.98 | 12.71 | 12.81 | 1.47 | -0.05 | 0.03 | 0.09 | *OsIDI1* |
| Os06g0486800 | 12.03 | 14.02 | 11.26 | 11.40 | 13.49 | 13.78 | 12.98 | 13.21 | 1.98 | 0.15 | 0.29 | 0.23 | *OsFDH* |
| Os09g0453800 | 13.64 | 16.06 | 13.05 | 13.19 | 13.89 | 13.92 | 13.90 | 13.98 | 2.41 | 0.14 | 0.03 | 0.08 | *OsIDI4* |
| Os03g0307300 | 14.01 | 18.21 | 2.60 | 2.77 | 5.07 | 13.62 | 4.43 | 7.67 | 4.20 | 0.17 | 8.55 | 3.24 | *OsNAS1* |
| Os03g0307200 | 15.54 | 18.52 | 4.15 | 3.93 | 4.76 | 14.23 | 3.25 | 10.36 | 2.98 | -0.22 | 9.47 | 7.11 | *OsNAS2* |
| Os03g0237100 | 9.78 | 13.72 | 8.83 | 8.52 | 8.99 | 9.65 | 8.33 | 8.27 | 3.94 | -0.31 | 0.66 | -0.06 | *OsDMAS1* |
| **Other Fe deficiency responsive genes** |  |  |  |  |  |  |  |  |  |  |  |  |  |
| Os01g0952800 | 5.37 | 10.92 | 5.30 | 12.10 | 4.20 | 10.32 | 2.18 | 12.66 | 5.56 | 6.79 | 6.13 | 10.48 | *OsIRO2* |
| Os03g0379300 | 8.67 | 12.28 | 8.90 | 13.27 | 8.45 | 11.73 | 8.24 | 13.43 | 3.62 | 4.36 | 3.28 | 5.19 | *OsIRO3* |
| Os12g0282000 | 4.96 | 12.16 | 6.28 | 13.77 | 2.45 | 11.08 | 1.96 | 13.70 | 7.20 | 7.49 | 8.63 | 11.74 | *OsMIR* |
| Os01g0647200 | 10.59 | 15.10 | 9.77 | 16.59 | 12.48 | 17.63 | 8.14 | 18.48 | 4.51 | 6.82 | 5.16 | 10.34 | *OsIMA1* |
| Os03g0667300 | 6.55 | 11.25 | 7.39 | 13.25 | 5.15 | 9.21 | 6.02 | 11.63 | 4.71 | 5.86 | 4.06 | 5.61 | *OsIRT2* |
| Os02g0649900 | 2.75 | 8.74 | 2.10 | 8.10 | 2.11 | 7.71 | 5.48 | 8.42 | 5.99 | 5.99 | 5.60 | 2.94 | *OsYSL2* |
| Os07g0258400 | 8.13 | 13.24 | 8.25 | 14.15 | 7.05 | 13.00 | 5.27 | 14.69 | 5.11 | 5.90 | 5.96 | 9.42 | *OsNRAMP1* |

The log2 values are shown.

**Figure 4.**
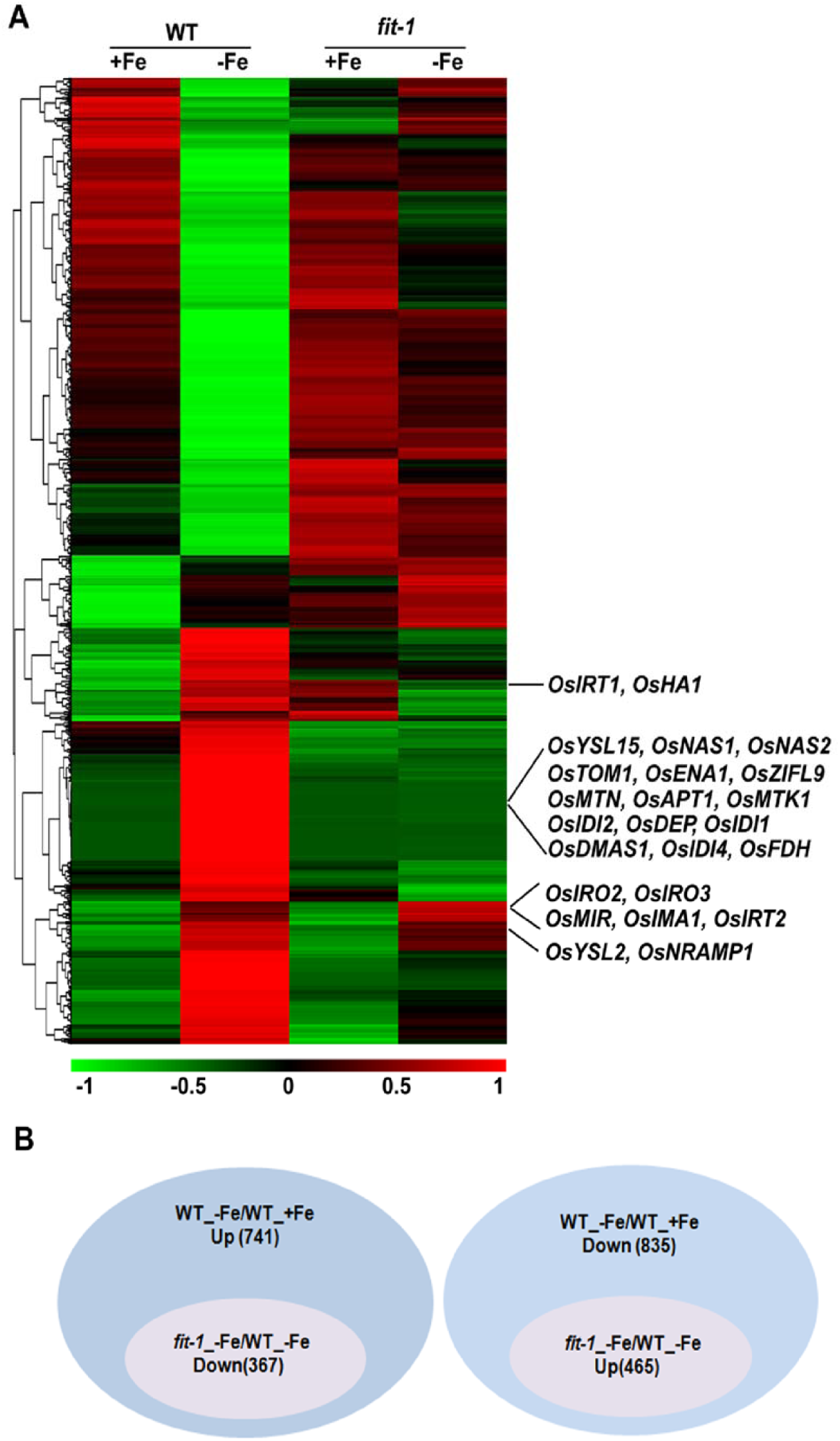
OsFIT transcriptional regulation of Fe homeostasis genes in the roots. **(A)** Heat map of 741 upregulated genes and 835 downregulated genes in wild type roots. **(B)** Fe deficiency responsive genes affected by OsFIT in the roots. Among the 741 upregulated genes by –Fe in the wild-type, 367 genes were down-regulated in the *fit-1* compared with in the wild type under –Fe. Among the 835 down-regulated genes by –Fe in the wild-type, 465 genes were upregulated in the *fit-1* compared with in the wild type under –Fe.

### Overexpression of *OsFIT* promotes Fe accumulation and expression of Fe-uptake genes

Considering that loss-of-function mutations to *OsFIT* resulted in increased sensitivity to Fe deficiency and decreased Fe accumulation, we examined whether upregulated *OsFIT* expression leads to enhanced tolerance to Fe deficiency and increased Fe accumulation. Thus, we generated *OsFIT-*overexpressing plants, among which two independent transgenic plants with high *OsFIT* expression levels were selected for subsequent analyses (Supplemental Figure S5A). There were no obvious differences between the *OsFIT-*overexpressing plants and the wild-type plants after a short-term (9 days) −Fe treatment (Supplemental Figure S5B). However, after a long-term (18 days) exposure to Fe deficiency, the older leaves of the *OsFIT-*overexpressing plants developed obvious rust spots, which were absent in the wild-type plants (Figure 5A, B). A subsequent comparison of the metal concentrations revealed that the *OsFIT-*overexpressing plants accumulated significantly more Fe and Zn, but not Cu, than the wild-type plants under both +Fe and –Fe conditions (Figure 5C; Supplemental Figure S5C). The rust spots on the leaves might result from the elevated Fe and Zn accumulation in the *OsFIT* overexpression lines. Next, we examined the expression of some Fe-deficiency-responsive genes that were downregulated in the *fit-1* mutant. The expression levels of all of the examined genes (*OsNAS1, OsNAS2, OsNAAT1, OsTOM1, OsYSL15*, and *OsIRT1*) increased in the *OsFIT-*overexpressing plants (Figure 5D). Our data imply that OsFIT positively regulates the expression of Fe-uptake-associated genes and promotes Fe uptake in rice.

**Figure 5.**
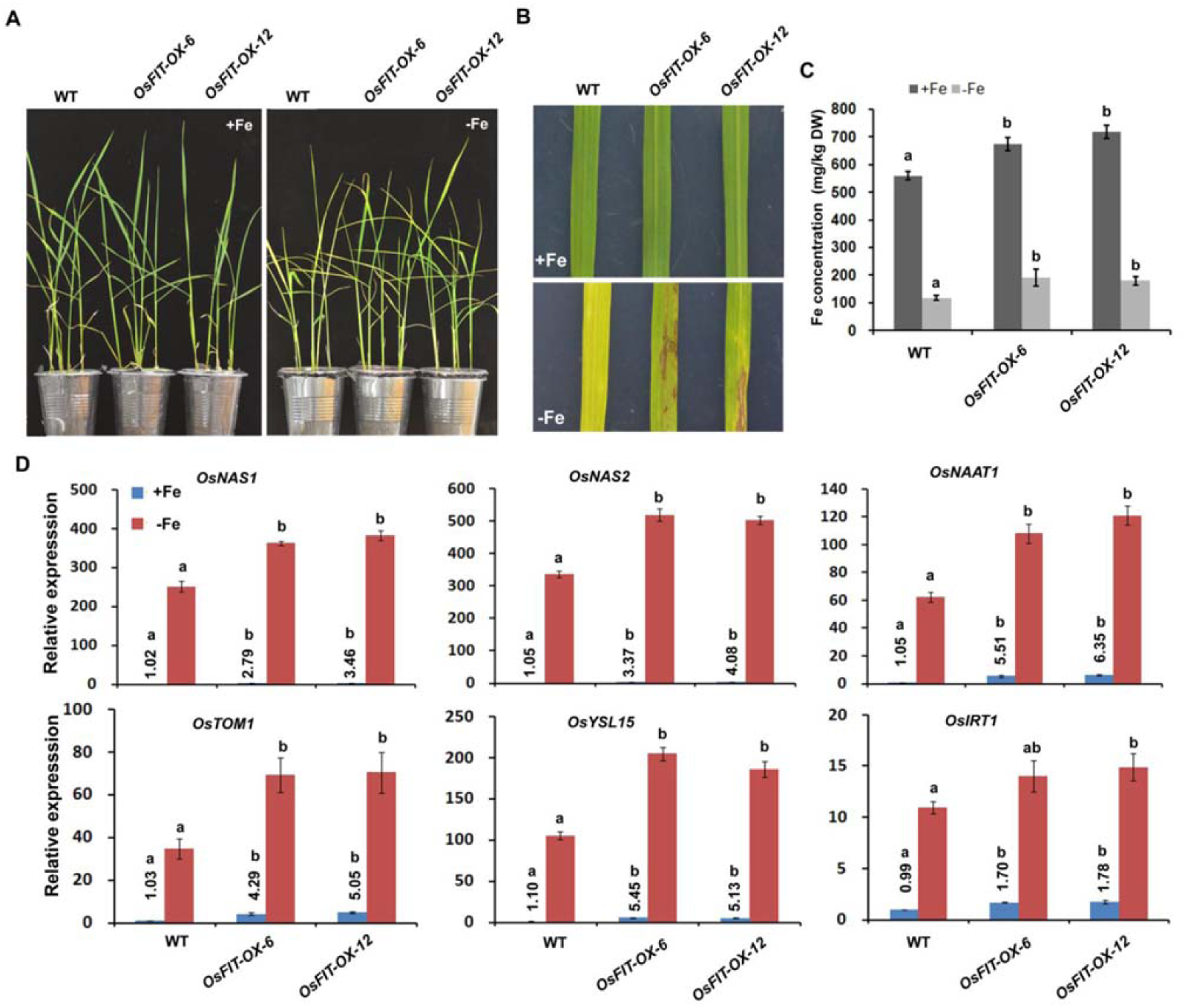
Analysis of *OsFIT* overexpression plants. **(A)** Growth of *OsFIT* overexpression plants. For +Fe growth, four-day-old seedlings germinated on wet paper were shifted in +Fe solution for 19 days. For –Fe growth, four-day-old seedlings germinated on wet paper were shifted in +Fe solution for 1 day and transferred to –Fe for 18 days. **(B)** The third leaves of seedlings in (A). **(C)** Fe concentration of shoots grown in +Fe solution in (A). Data represent means ± SD (*n* = 3). Ddifferent letters above each bar indicate statistically significant differences as determined by one-way ANOVA followed by Tukey’s multiple comparison test (P < 0.05). **(D)** Expression of Fe uptake genes. Four-day-old seedlings germinated on wet paper were grown in +Fe for 5 days and then transferred to +Fe or –Fe for 5 days. Roots were used for RNA extraction. Data represent means ± SD (*n* = 3). Different letters above each bar indicate statistically significant differences as determined by one-way ANOVA followed by Tukey’s multiple comparison test (P < 0.05).

### Genetic relationship between *OsFIT* and *OsIRO2*

Although we confirmed that OsFIT interacts with OsIRO2 (Figure 1), it remained unclear how these two transcription factors regulate Fe homeostasis. We observed that the *OsIRO2* transcript levels increased in the *fit-1* mutant (Table 1), implying that *OsIRO2* expression is not positively regulated by OsFIT. To determine whether *OsFIT* is positively regulated by OsIRO2, we examined its expression in the *iro2-1* mutant (Zhang et al., 2019). Interestingly, the *OsFIT* transcript level increased in the *iro2-1* mutant (Supplemental Figure S6). To investigate the genetic relationship between *OsFIT* and *OsIRO2*, we generated the *fit-1 iro2-1* double mutant by crossing two single mutants. When grown in the +Fe solution, the single and double mutants were phenotypically similar to the wild-type control (Figure 6A). When grown in the −Fe solution, the single and double mutants, but not the wild-type control, developed chlorotic leaves; however, there were no observable differences between the single and double mutants. Moreover, the Fe concentration of the *fit-1 iro2-1* double mutant plants was as low as that of the single mutants *fit-1* and *iro2-1* (Figure 6B). We subsequently determined the expression of Fe-deficiency-responsive genes in the wild-type plants as well as in the single and double mutants (Figure 6C). The downstream Fe-uptake genes were expressed at a similar level between the single and double mutants. Collectively, our data suggest that OsIRO2 and OsFIT function in the same node of the Fe homeostasis signaling network.

**Figure 6.**
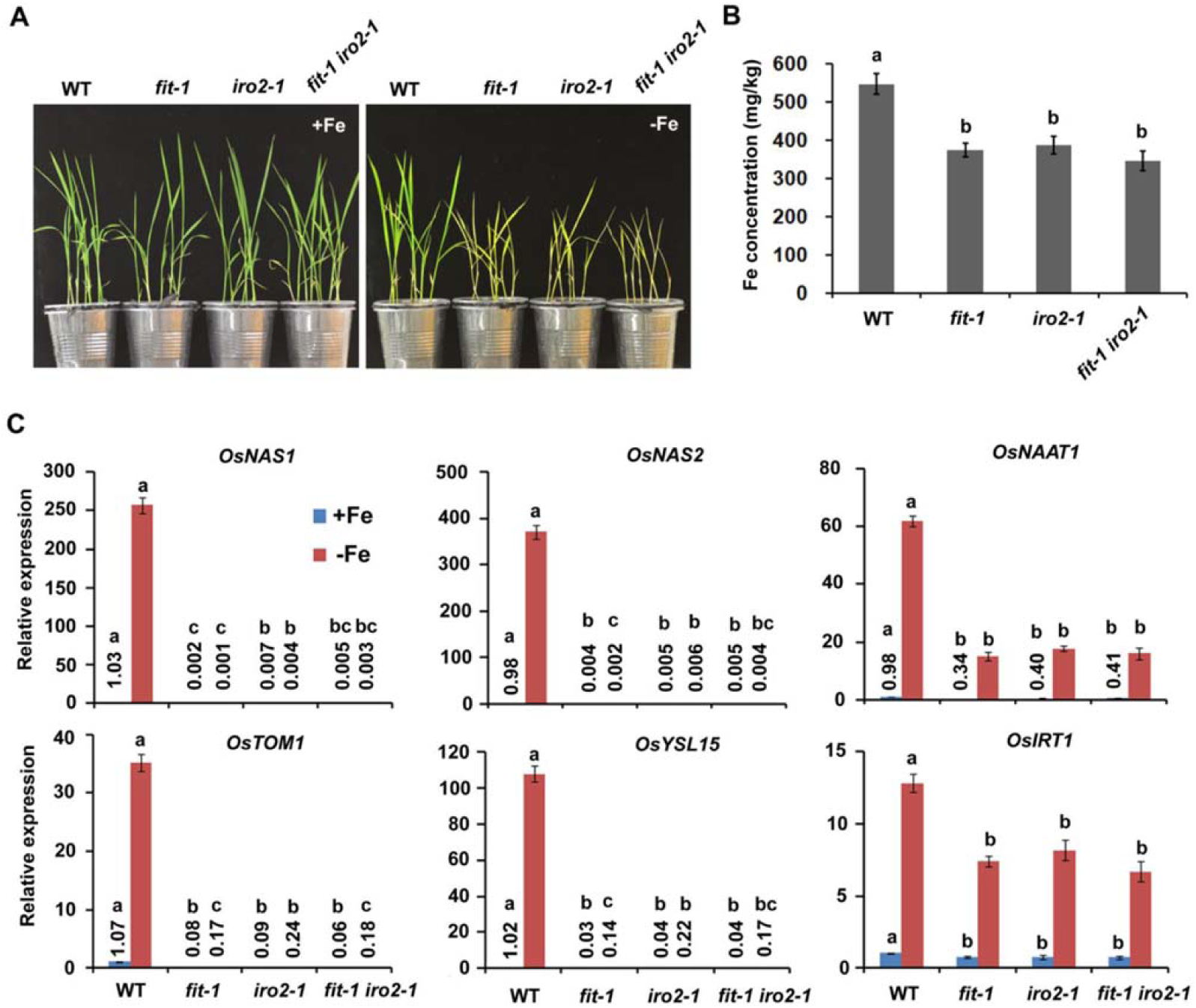
Genetic interaction between OsIRO2 and OsFIT. **(A)** Growth of various mutant seedlings. Seeds were germinated on wet paper for four days. For +Fe growth, four-day-old seedlings germinated on wet paper were shifted in +Fe for 10 days. For –Fe growth, four-day-old seedlings germinated on wet paper were shifted in +Fe for 1 day and transferred to –Fe for 9 days. **(B)** Fe concentration of shoots grown in +Fe solution in (A). Data represent means ± SD (*n* = 3). Different letters above each bar indicate statistically significant differences as determined by one-way ANOVA followed by Tukey’s multiple comparison test (P < 0.05). **(C)** Expression of Fe uptake genes. Four-day-old seedlings were grown in +Fe for 5 days and transferred to +Fe or –Fe for 5 days. Roots were used for RNA extraction. Data represent means ± SD (*n* = 3). Different letters above each bar indicate statistically significant differences as determined by one-way ANOVA followed by Tukey’s multiple comparison test (P < 0.05).

### OsFIT and OsIRO2 function as a transcription complex

Although OsFIT and OsIRO2 interact with each other and positively regulate Fe homeostasis, the molecular mechanism underlied is unclear. Both OsFIT and OsIRO2 are bHLH transcription factors and they regulate the expression of numerous Fe deficiency responsive genes. We proposed that some of these genes are the direct targets of OsIRO2 and OsFIT. Generally, bHLH transcription factors regulate their targets by binding to the E-boxes within the promoters (Fisher and Goding, 1992). A sequence analysis revealed several E-boxes within the promoters of Fe-deficiency-responsive genes (*OsNAS1, OsNAS2, OsNAAT1, OsDMAS1, OsTOM1*, and *OsYSL15*) (Supplemental Figure S7A). To determine whether OsFIT and OsIRO2 bind to the E-boxes, we performed EMSAs with *OsNAS2* and *OsYSL15* promoter probes. The full-length *OsFIT* and *OsIRO2* were respectively fused with the glutathione *S*-transferase (GST) and the recombinant proteins GST-OsFIT and GST-OsIRO2 were expressed in and purified from *E. coli* cells. We observed that the biotin probe was able to bind to GST-OsIRO2, but not GST-OsFIT or GST alone (Figure 7A). The binding of GST-OsIRO2 by the biotin probe was inhibited by the addition of increasing amounts of the unlabeled probes (cold probe), but not the mutated probe (cold probe-m). These observations suggest that OsIRO2 can bind to the *OsNAS2* and *OsYSL15* promoters. Considering that OsFIT interacts with OsIRO2, we assessed whether OsFIT affects the binding of OsIRO2 to DNA. Specifically, both OsFIT and OsIRO2 were incubated with the probes. The presence of OsFIT appeared to enhance the binding of OsIRO2 to the probes.

**Figure 7.**
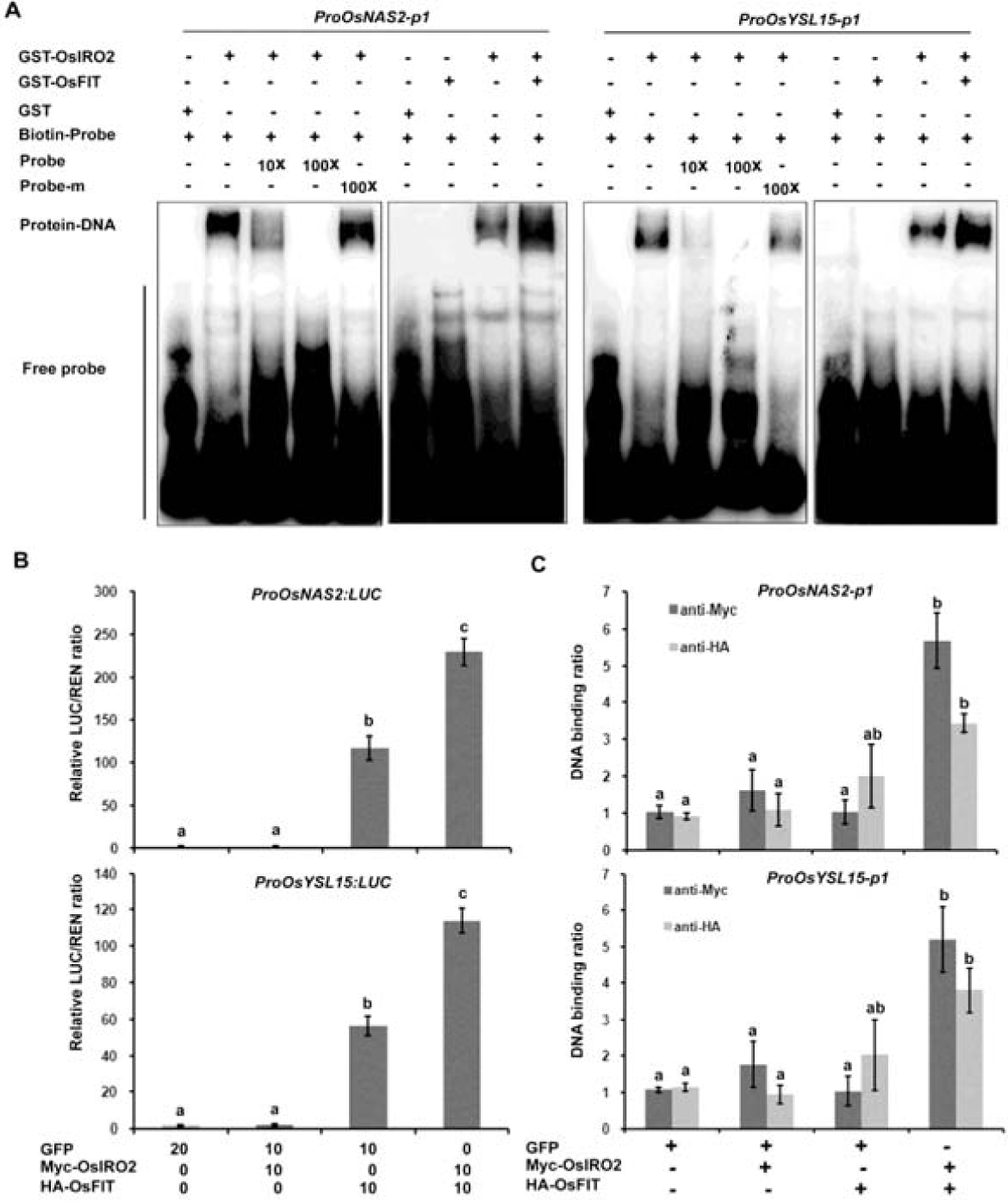
OsIRO2 and OsFIT mutually regulate the expression of *OsNAS2* and *OsYSL15*. **(A)** EMSA showing that OsIRO2 directly binds the *OsNAS2* and *OsYSL15* promoters. GST-OsIRO2 and/or OsFIT were incubated with the biotin-labeled probes. Biotin-probe, biotin-labeled probe; cold-probe, unlabeled probe; cold-probe-m, unlabeled mutated probe. Biotin probe incubated with GST served as the negative control. **(B)** Activation of target genes by OsIRO2 and OsFIT. The LUC/REN ratio represents the LUC activity relative to the internal control (REN driven by the 35S promoter). The numbers 0, 10 and 20 indicate that 0, 10, and 20 μg of plasmid were used for the corresponding vectors in the transient expression assay. Data represent means ± SD (*n* = 3). Different letters above each bar indicate statistically significant differences as determined by one-way ANOVA followed by Tukey’s multiple comparison test (P < 0.05). **(C)** ChIP-qPCR analysis. The numbers in the parenthesis indicate the relative proportion of agrobacteria in each combination. Data represent means ± SD (*n* = 3). Different letters above each bar indicate statistically significant differences as determined by one-way ANOVA followed by Tukey’s multiple comparison test (P < 0.05).

We then examined the transactivation ability of OsIRO2 and OsFIT by the transient expression assay involving a LUC-based effector-reporter system in the wild-type rice protoplasts (Figure 7B). The *OsNAS2* and *OsYSL15* promoters were fused to the *LUC* gene as the reporters (*ProOsNAS2:LUC* and *ProOsYSL15:LUC*). The *Myc-OsIRO2* and *HA-OsFIT* under the control of the CaMV 35S promoter functioned as the effectors. When the effector *Myc-OsIRO2* was co-expressed with the reporters, the LUC activity did not increase significantly. In contrast, the effector *HA-OsFIT* substantially increased the LUC activity of both reporters. When both effectors were expressed simultaneously, the LUC activity was higher than that induced by the *HA-OsFIT* effector alone. These results suggest that OsFIT, but not OsIRO2, activates the expression of their target genes.

To further investigate whether OsFIT and OsIRO2 functionally depend each other, Myc-OsIRO2 and the reporters were coexpressed in the *fit-1* protoplasts, and HA-OsFIT and the reporters in the *iro2-1* protoplasts. As a result, the LUC activity of reporters did not increase (Supplemental Figure S7B), suggesting that both OsIRO2 and OsFIT are required for the expression of their target genes.

Subsequently, we conducted ChIP-qPCR assays to investigate the binding of OsIRO2 and OsFIT to their target genes *in vivo.* When Myc-OsIRO2 or HA-OsFIT was expressed with the effectors, no significant binding to target sequences was detected. In contrast, when Myc-OsIRO2 and HA-OsFIT were co-expressed, they could bind to the promoters of *OsNAS2* and *OsYSL15* (Figure 7C). These results further suggest that OsFIT and OsIRO2 function as a transcription complex to regulate the expression of their target genes.

### *OsFIT* expression is positively regulated by OsPRI1, OsPRI2, and OsPRI3

Although OsFIT and OsIRO2 positively regulate the expression of many Fe-deficiency-responsive genes, the transcription of *OsFIT* and *OsIRO2* is also induced by Fe deficiency. We previously revealed that OsPRI1, OsPRI2, and OsPRI3 directly activate *OsIRO2* expression (Zhang et al., 2017, 2019). Therefore, we evaluated whether *OsFIT* expression is also positively regulated by OsPRI1, OsPRI2, and OsPRI3. An analysis of *OsFIT* indicated that the its expression levels decreased in the *pri1-1, pri2-1*, and *pri3-1* mutant plants and increased in the *OsPRI2*- and *OsPRI3*-overexpressing plants (Figure 8). These data suggest that OsPRI1, OsPRI2, and OsPRI3 stimulate *OsFIT* expression under Fe-deficient conditions.

**Figure 8.**
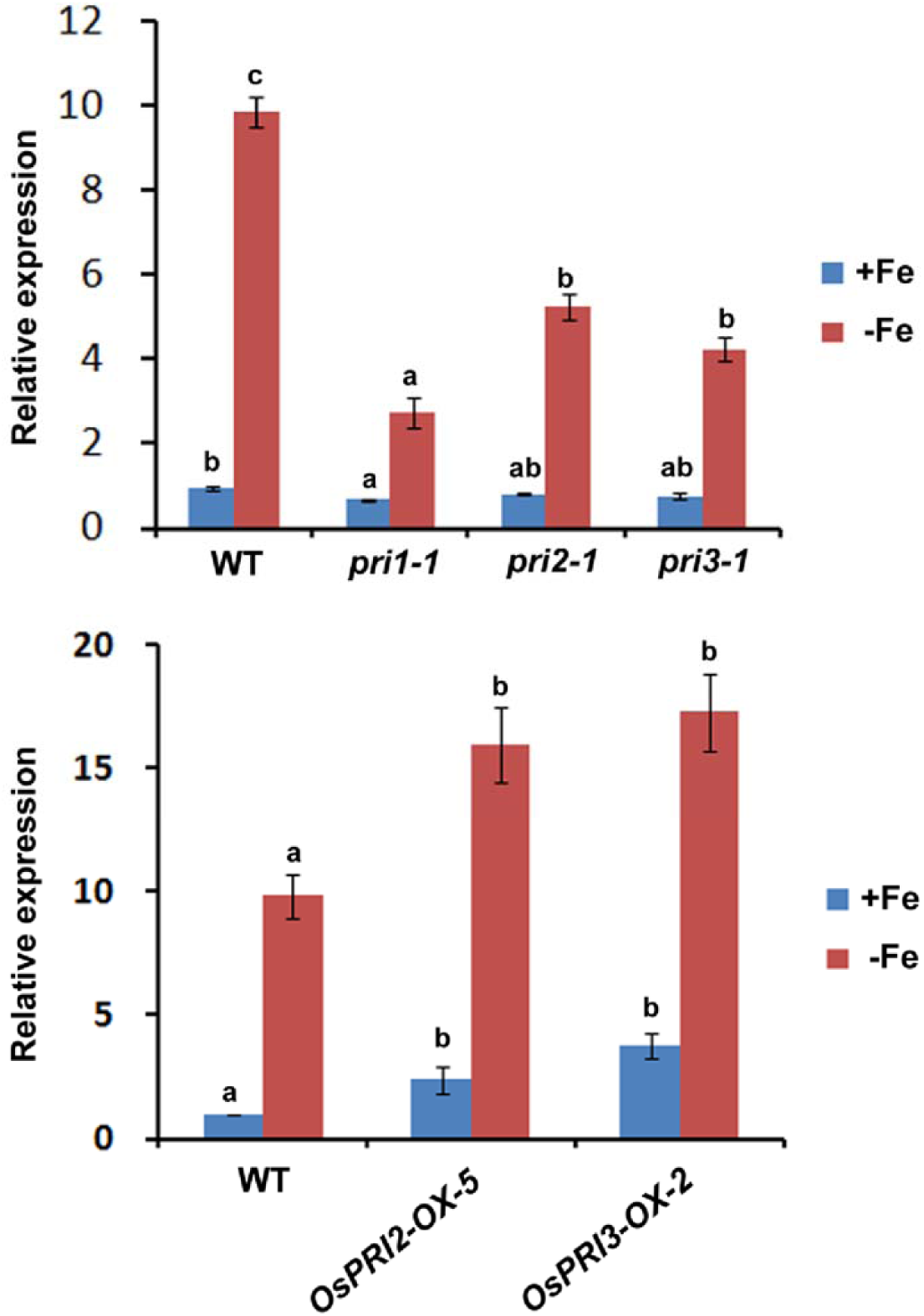
*OsFIT* is regulated positively by the OsPRI proteins. Four-day-old seedlings germinated on wet paper were grown in +Fe for 5 days and transferred to +Fe or –Fe for 5 days. Roots were used for RNA extraction. Data represent means ±SD (*n* = 3). Different letters above each bar indicate statistically significant differences as determined by one-way ANOVA followed 865 by Tukey’s multiple comparison test (P < 0.05).

## DISCUSSION

In response to Fe deficiency, plants modify the expression of numerous genes to maintain Fe homeostasis. However, the signal transduction network regulating the expression of Fe-homeostasis-associated genes has not been comprehensively characterized. As a key component of the Fe homeostasis signaling network, OsIRO2 positively regulates rice Fe homeostasis. Here, we identified and characterized its interaction partner OsFIT. When this manuscript was under review, Wang et al. (2020) reported the similar functions of OsFIT/OsbHLH156 in rice Fe homeostasis. In this study, we further revealed that OsFIT and OsIRO2 function in the same genetic node and form a transcription complex to regulate rice Fe homeostasis.

### Similarities and differences between OsFIT and AtFIT

Earlier investigations proved that AtFIT interacts with Arabidopsis bHLH Ib TFs and positively regulates the strategy I system in Arabidopsis (Yuan et al., 2008; Wang et al., 2013). Another study indicated OsIRO2 is a rice ortholog of Arabidopsis bHLH Ib TFs (Ogo et al., 2007). Because of the functional redundancy among Arabidopsis bHLH Ib TFs, their single or double mutants have no visible Fe-deficiency symptoms and their triple mutants exhibit Fe-deficiency symptoms that are not as extensive as those of the *fit* mutant (Wang et al., 2013). In contrast, we observed that the Fe-deficiency symptoms of the single *iro2-1* mutant were as strong as those of the *fit* mutants (Figure 6A, B). These results suggest that OsIRO2 is similar to Arabidopsis bHLH Ib TFs. Moreover, OsIRO2 may be the only rice ortholog of Arabidopsis bHLH Ib TFs. Similar to the interaction between AtFIT and bHLH Ib, OsFIT interacts with OsIRO2 and positively regulates the expression of Fe-uptake genes in rice (Table 1; Figure 5D, 6C). The expression of *AtFIT* is induced by Fe deficiency and is positively regulated by bHLH IVc TFs (Zhang et al., 2015; Li et al., 2016; Liang et al., 2015). Similarly, *OsFIT* expression is induced by Fe deficiency via OsPRI1, OsPRI2, and OsPRI3 (Figure 8), which are orthologs of Arabidopsis bHLH IVc. Furthermore, the OsFIT protein sequence is 33.68% similar to the AtFIT sequence (Supplemental Figure S1B). Given these similarities between OsFIT and AtFIT, we assume that OsFIT is a rice ortholog of AtFIT.

The following evidence indicates the differences of physiological functions of OsFIT and AtFIT: (1) *AtFIT* is specifically expressed in the roots, whereas *OsFIT* is expressed and functional in the roots and leaves (see discussion below); (2) *AtFIT* overexpression has no significant effect on the expression of its downstream genes *AtIRT1* and *AtFRO2* (Colangelo and Guerinot, 2004), but *OsFIT* overexpression activates its downstream Fe-uptake-associated genes (Figure 5D); and (3) AtFIT regulates strategy I in Arabidopsis, whereas OsFIT regulates both strategy I and II in rice (see discussion below).

### OsFIT is involved in both strategy I and II

Rice plants possess the strategy II Fe-uptake system specific to graminaceous plants, but also a partial strategy I Fe-uptake system, which is advantageous for growth in submerged conditions. Ishimaru et al. (2006) confirmed that rice takes up both Fe(III)-phytosiderophore and Fe(II). The results of a recent study suggest that rice uses strategy II to absorb Fe under Fe-deficient conditions, whereas strategy I is applied under Fe-sufficient conditions (Liu et al., 2019).

Rice strategy II-associated genes include *OsNAS1, OsNAS2, OsDMAS1, OsNAAT1, OsTOM1*, and *OsYSL15*. The expression levels of all of the analyzed strategy II-associated genes were considerably downregulated in the *fit* mutants (Table 1; Figure 6C), suggesting the strategy II Fe-uptake system was impaired. Correspondingly, the Fe concentration was lower in the *fit* mutants than in the wild-type control when Fe(III) was the only Fe source (Figure 3E). These data suggest that OsFIT is a crucial regulator of the strategy II Fe-uptake system.

It is unclear which genes are responsible for the strategy I system in rice. In Arabidopsis, AtIRT1 is a key component of strategy I. Although the AtIRT2 and AtIRT1 amino acid sequences are similar, AtIRT2 cannot rescue the phenotypes caused by loss-of-function mutations to *AtIRT1*. Unlike AtIRT1 which is a plasma-membrane protein (Vert et al., 2002), AtIRT2 is localized to intracellular vesicles, and hence may be responsible for compartmentalization of iron (Vert et al., 2009). A previous study revealed a lack of significant differences between the loss-of-function *irt2-1* mutant and the wild-type plants (Varotto et al., 2002), implying that AtIRT2 is not a key component of the strategy I system. Rice contains OsIRT1 and OsIRT2, which are the counterparts of AtIRT1 and AtIRT2, respectively. However, unlike AtIRT2, OsIRT2 is a plasma-membrane protein (Ishimaru et al., 2006). Thus, it is still unclear which one of OsIRT1 and OsIRT2 is responsible for the uptake of Fe(II) in rice. In the current study, *OsIRT1* expression was downregulated in the *fit* mutants (Table 1; Figure 6C), implying that OsFIT positively regulates *OsIRT1* expression. In contrast, *OsIRT2* was upregulated in the *fit-1* mutant (Table 1). Similarly, *AtIRT1* is also downregulated in the Arabidopsis *fit* mutant (Colangelo and Guerinot, 2004). Thus, it is very likely that OsIRT1 functions in the translocation of Fe(II) from soil to roots. Rhizosphere acidification mediated by plasma membrane H^+^-ATPases is a key step in the strategy I Fe-uptake system. The expression of *OsHA1* (Os05g0319800), which encodes a plasma membrane H^+^-ATPase, is induced in response to Fe deficiency, indicating that OsHA1 may be a component of the rice strategy I Fe-uptake system. The transcription of *OsHA1* decreased significantly in the *fit-1* mutant (Table 1), suggesting *OsHA1* expression is positively regulated by OsFIT.

An earlier study on the *naat1* mutant indicates that the strategy I system is necessary for rice homeostasis (Cheng et al., 2007). The *naat1* mutant, in which the strategy II system is damaged, cannot survive a nutrient solution with Fe(III) as the only Fe source. Additionally, this mutant activates the strategy I system and accumulates more Fe than the wild-type control in the Fe(II) solution. However, the *fit* mutants with a severely inhibited strategy II system did not activate strategy I. In fact, less Fe accumulated in the *fit* mutants than in the wild-type plants when grown in the Fe(II) solution (Figure 3E), implying the strategy I system was slightly inhibited. Therefore, we propose that OsFIT regulates both strategy I and II.

### The OsFIT-OsIRO2 complex regulates Fe homeostasis

The *OsIRO2* gene is expressed in the root and leaf vascular tissues under Fe-sufficient conditions, but its expression can extend to all root and leaf tissues under Fe-deficient conditions (Ogo et al., 2011). In addition to controlling the Fe-uptake genes in the roots, OsFIT also mediates the expression of some Fe-uptake genes in the shoots. The expression levels of several Fe-uptake genes, such as *OsNAS1/2, OsYSL15, OsENA1, OsTOM1*, and *OsZIFL9*, which are upregulated in wild-type shoots under Fe-deficient conditions (Table 1), are downregulated in the *fit-1* shoots, indicating OsFIT is functional in the shoots. The induction of *OsNAS1* and *OsNAS2* expression is completely blocked in the *fit-1* roots, but not in the *fit-1* shoots. It is likely that other transcription factors are also involved in the activation of *OsNAS1* and *OsNAS2* expression in the shoots.

Although rice and Arabidopsis evolved different Fe-uptake strategies, the expression of Fe-uptake genes in both species is controlled by a very similar regulatory network. In Arabidopsis, AtFIT and bHLH Ib TFs function downstream of bHLH IVc TFs (Zhang et al., 2015; Li et al., 2016; Liang et al., 2017). Similarly, *OsFIT* and *OsIRO2* expression levels are positively regulated by OsPRI1, OsPRI2, and OsPRI3 (Figure 8). Although *OsFIT* transcript levels were elevated in the *iro2-1* mutants (Supplemental Figure S6), these mutants still exhibited Fe-deficiency symptoms. Similarly, *OsIRO2* was also highly expressed in the *fit-1* mutant (Table 1). These observations suggest that the elevated *OsFIT* is not sufficient to rescue the loss-of-function of OsIRO2, and vice versa. Meanwhile, the deficiency symptoms and the expression of Fe-uptake genes in the *fit-1 iro2-1* double mutants were similar to those of the single mutants *fit-1* and *iro2-1*, implying that OsFIT and OsIRO2 function in the same genetic node (Figure 6).

Our results indicate that OsFIT physically interacts with OsIRO2 (Figure 1). OsIRO2 is preferentially expressed in the cytoplasm, and the upregulated expression of OsFIT results in the increased nuclear accumulation of OsIRO2 (Figure 2C). A similar result was also reported by Wang et al. (2020). A latest study found that AtFIT promotes the nuclear accumulation of AtbHLH39, an Arabidopsis homolog of OsIRO2, in Arabidopsis (Trofimov et al., 2019). Similarly, Arabidopsis bHLH IVc TFs promote the nuclear accumulation of AtbHLH121 which is required for the maintenance of Arabidopsis Fe homeostasis (Kim et al., 2019; Gao et al., 2020; Lei et al., 2020). Although we did not observe the localization change of OsIRO2 in response to Fe deficiency in our transient expression assays, the increased nuclear accumulation of OsIRO2 under Fe deficiency condition was observed by Wang et al. (2020). Further investigations are required to reveal the biological relevance of OsIRO2 localization change with the Fe deficiency response.

Although OsIRO2 binds to the target promoters and OsFIT may enhance its binding activity (Figure 7A), OsIRO2 alone could not activate the expression of the target genes (Figure 7B). Therefore, OsIRO2 may lack a transcription activation ability. In contrast, OsFIT did not bind to the promoters of *OsNAS2* and *OsYSL15* in our EMSA, but OsFIT activates the latter (Figure 7A, B). It is plausible that OsFIT lacks a DNA binding ability. It is noteworthy that OsIRO2 binds to its target promoters *in vivo* only in the presence of OsFIT (Figure 7C), agreeing with that OsFIT promotes the accumulation of OsIRO2 in the nucleus where it binds to the target DNA. Meanwhile, it is also noted that the activation of the *OsNAS2* and *OsYSL15* promoters by OsFIT occurs in the wild-type (Figure 7B), but not in the *iro2-1* mutant (Supplemental Figure S7B), implying that the function of OsFIT may depend on the DNA binding ability of OsIRO2. Therefore, we propose that both OsFIT and OsIRO2 are required for the expression of their downstream genes; and they interact with each other to form a functional transcription complex to regulate Fe-uptake associated genes.

### A working model of OsFIT and OsIRO2

OsHRZ1 is a putative Fe-binding sensor which negatively regulates Fe homeostasis in rice (Kobayashi et al., 2013). Our recent work revealed that OsHRZ1 interacts with and degrades OsPRI1, OsPRI2 and OsPRI3 which directly activate the expression of *OsIRO2* and *OsIRO3* (Zhang et al., 2017, 2019). OsIRO2 positively and OsIRO3 negatively regulate Fe homeostasis respectively (Ogo et al., 2007; Zheng et al., 2010). Here, we further revealed that *OsFIT* is positively regulated by OsPRI1/2/3, and OsFIT and OsIRO2 form a transcription complex to regulate Fe homeostasis. A working model of OsIRO2 and OsFIT was developed (Figure 9). Specifically, when plants are exposed to Fe-deficient conditions, OsPRI1, OsPRI2 and OsPRI3 activate the expression of *OsIRO2* and *OsFIT.* Under Fe sufficient conditions, OsIRO2 is preferentially located in the cytoplasm. Under Fe deficienct conditions, OsFIT promotes the accumulation of OsIRO2 in the nucleus, and then OsIRO2 and OsFIT form a functional transcription complex to activate the expression of Fe-uptake genes.

**Figure 9.**
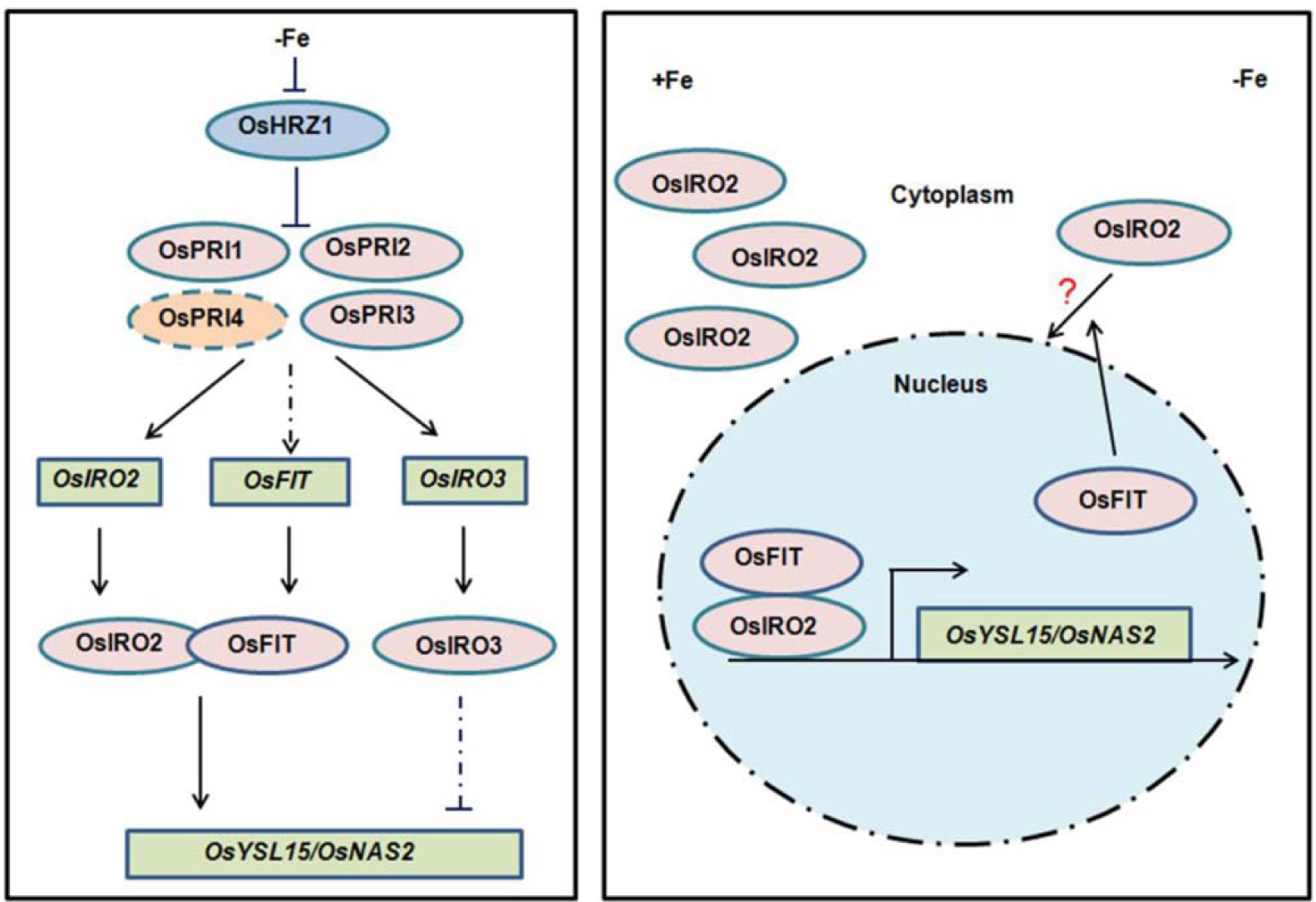
A proposed working model of OsFIT and OsIRO2. Fe deficiency response pathways in rice. OsHRZ1 interacts with and inhibits OsPRI1, OsPRI2 and OsPRI3 which activate the expression of *OsIRO2, OsFIT* and *OsIRO3*. OsPRI4, a paralog of OsPRI1, OsPRI2 and OsPRI3, may play a redundant role in Fe homeostasis. OsIRO2 and OsFIT form a heterodimer to initiate the expression of their downstream target genes. OsIRO3 negatively regulates the expression of Fe deficiency responsive genes. OsIRO2 is preferentially expressed in the cytoplasm. OsFIT is specifically expressed in the nucleus. OsFIT facilitates the nuclear accumulation of OsIRO2 in the nucleus. OsFIT and OsIRO2 function as a transcription complex to control the expression of Fe-uptake genes.

## MATERIALS AND METHODS

### Plant materials and growth conditions

Rice (*Oryza sativa* L. cv. Nipponbare) seeds were germinated in wet paper for five days and moved to hydroponic culture or soils. For hydroponic culture assays, half-strength Murashige and Skoog (MS) media (pH5.6-5.8) with 0.1 mM Fe(III)-EDTA or 0.1 mM Fe(II)-EDTA or without Fe. For Fe(II) solution, 0.1 mM hydroxylamine was supplemented to avoid the oxidation of Fe(II). The nutrient solution was exchanged every 3 days. For soil assays, plants were grown in waterlogged or aerobic soil. The plants were watered with tap water. Plants were grown in a growth chamber at 28°C/20°C (day/night) and 60 % to 70 % humidity, bulb type light with a photon density of ∼300 μmol m^-2^ s^-1^ and a photoperiod of 14 h.

### Gene expression analysis

Total RNA from rice roots or shoots was reverse transcribed using an oligo dT and HiScript II Q RT SuperMix for qPCR (+gDNA wiper) (Vazyme, China) following the manufacturer’s protocol. qPCR was performed on a Light-Cycler 480 real-time PCR machine (Roche, Switzerland) by the use of an AceQ Universal SYBR qPCR Master Mix (Vazyme, China). All PCR amplifications were performed in triplicate, with the Os*ACTIN1* gene as an internal control. Primers used for qPCR are listed in Supplemental Table S2.

### Fe measurement

The harvested plants were rinsed with distilled water and then blotted using paper towels. The shoots were then separated and dried at 65 °C for one week. For each sample, about 500 mg dry weight of shoots were digested with 5 ml of 11 M HNO_3_ and 2 ml of 12 M HClO_4_ for 30min at 220°C. Metal concentrations were measured using Inductively Coupled Plasma Mass Spectrometry (ICP-MS).

### Yeast two-hybrid assays

The yeast two-hybrid assays were carried out according to the manufacturer’s protocol. The OsIRO2 N-terminal fragment (aa 1-150) was subcloned to the pGBKT7 plasmid as bait. Yeast transformation was performed according to the Yeastmaker Yeast Transformation System 2 User Manual (Clontech). More than 1 × 10^6^ yeast clones were screened in synthetic defined (SD) medium minus Trp, Leu, His, and Ade. The plasmids of the positive clones were extracted and then retransformed into yeast for the double check on selective SD plates. Positive clones were selected for sequencing.

In yeast two-hybrid assays, the full-length OsFIT was subcloned to pGADT7 and then co-transformed with pGBKT7-OsIRO2-N. Yeast transformation was performed according to the Yeastmaker Yeast Transformation System 2 User Manual (Clontech). The primers used are listed in Supplemental Table S2.

### Generation of Plasmids Used for Transgenic Plants

To ensure the gene targeting efficiency and avoid off-targets, target sites were designed by the use of CRISPR-GE (Xie et al., 2017). The editing vectors were constructed as described previously (Liang et al., 2016). Briefly, the OsU6a promoter driving the sgRNA containing a single target site was cloned into the pMH-SA vector by the restriction enzyme sites *Spe*I and *Asc*I (Liang et al., 2016).

For the construction of overexpression vector, the HA-OsFIT fusion sequence was obtained from the GAD-OsFIT vector and cloned between the maize ubiquitin promoter and the NOS terminator in the pUN1301 binary vector.

For the construction of *ProOsFIT:GUS* vector, the 3.2kb sequence upstream of *OsFIT* was subcloned into pCAMBIA1300-GUS plasmid with the *Sac* I site by a modified Gibson Assembly method (Zhu et al., 2014). Histochemical GUS staining assays were performed by the use of GUS histochemical assays kit (Real-Times, China) following the manufacturer’s protocol.

### Tripartite split-GFP assays

*Agrobacterium tumefaciens* strain EHA105 was used in the transient expression experiments. The sfGFP was divided into three parts, GFP1-9, GFP10 and GFP11, as described previously (Liu et al., 2018) and then they were subcloned into the pER8 vector to generate pTG1-9, pTG10 and pTG11 respectively. The pTG10 plasmid was linearized by the *Xho* I and fused in frame with OsIRO2 by a modified Gibson Assembly method (Zhu et al., 2014). Similarly, the pTG11 plasmid was linearized by the *Xho* I and fused in frame with OsFIT. Agrobacteria were incubated in LB liquid media. When growth reached an OD600 of approximately 3.0, the bacteria were spun down gently (3200 g, 5 min), and the pellets were resuspended in infiltration buffer (10 mM MgCl_2_, 10 mM MES, pH 5.6) at a final OD600 of 1.5. A final concentration of mM acetosyringone was added, and the bacteria were kept at room temperature for at least 2 h without shaking. For coinfiltration, the combination of different constructs as indicated in Figure 1B were mixed prior to infiltration. Leaf infiltration was conducted in 3-week-old *Nicotiana benthamiana*. The abaxial sides of leaves were injected with 20 μM β-estradiol 24 h before observation.

### Co-IP Assays

Agrobacterium strain EH105 cells carrying the *Pro35S:HA-GFP, Pro35S:Myc-OsIRO2*, or *Pro35S:HA-OsFIT* constructs were combined as indicted and transiently infiltrated into *N. benthamiana* leaves. The plants were grown in dark for 2 d and lysed. The extracts were incubated with 5 μL anti-HA antibodies coupled with 30 μL Protein-A Sepharose (GE Healthcare) overnight at 4 °C. The Sepharose was washed three times with protein extraction buffer. The samples were analyzed by immunoblotting using an anti-Myc antibody.

### Subcellular Localization

The full-length OsFIT was fused with GFP to generate OsFIT-GFP and the full-length OsIRO2 with mCherry to generate OsIRO2-mCherry. The plasmids above were transformed into agrobacteria. Agrobacteria were incubated in LB liquid media. When growth reached an OD600 of approximately 3.0, the bacteria were spun down gently (3200 g, 5 min), and the pellets were resuspended in infiltration buffer (10 mM MgCl_2_, 10 mM MES, pH 5.6) at a final OD600 of 1.0. Agrobacteria were mixed at a ratio indicated in Figure 2C and a final concentration of 0.2 mM acetosyringone was added. The agrobacteria were kept at room temperature for at least 2 h without shaking. Leaf infiltration was conducted in 3-week-old *N. benthamiana*. Excitation laser wave lengths of 488 nm and 563 nm were used for imaging GFP and mCherry signals, respectively. Total intensities of the nucleus and the cytoplasm fluorescence were measured separately by Image J. The ratio was calculated for each individual cell.

### Transient Luciferase Expression Assay

GFP, Myc-OsIRO2 and HA-OsFIT were subcloned to pGreenII 62SK as the effectors, and the *OsNAS2* and *OsYSL15* promoters were subcloned respectively to pGreen0800-LUC as the reporters (*ProOsNAS2:LUC* and *ProOsYSL15:LUC*).

Rice mesophyll cell protoplasts were prepared as described previously (Zhang et al., 2011). Green tissues from the stem and sheath of ten-day-old seedlings grown on half MS medium were used. A bundle of rice plants (about 30 seedlings) were cut together into approximately 0.5 mm strips using a sharp razor. The strips were immediately transferred into 0.6 M mannitol for 10 min in the dark. After discarding the mannitol, the strips were incubated in an enzyme solution (1.5% Cellulase RS, 0.75% Macerozyme R-10, 0.6 M mannitol, 10 mM MES at pH 5.7, 10 mM CaCl_2_ and 0.1% BSA) for 4-5 h in the dark with gentle shaking (60-80rpm). After the enzymatic digestion, an equal volume of W5 solution (154 mM NaCl, 125 mM CaCl_2_, 5mM KCl and 2 mM MES at pH 5.7) was added, followed by vigorous shaking by hand for 10 sec. Protoplasts were released by filtering through 40 nylon meshes into round bottom tubes with 3-5 times wash of the strips using W5 solution. The pellets were collected by centrifugation at 1,500 rpm for 3 min with a swinging bucket. After washing once with W5 solution, the pellets were then resuspended in MMG solution (0.4 M mannitol, 15 mM MgCl_2_ and 4 mM MES at pH 5.7) at a concentration of 2 × 10^6^ cells mL^-1^.

In transient luciferase expression assay, plasmids were transfected into protoplasts as described previously (Yoo et al., 2007). A Dual Luciferase kit (Promega) was used to detect reporter activity. The *Renilla luciferase* gene driven by the 35S promoter was used as an internal control.

### Chromatin Immunoprecipitation (ChIP) assays

The effector plasmids (pGreenII 62SK-GFP, pGreenII 62SK-Myc-OsIRO2, and pGreenII 62SK-HA-OsFIT) and the promoter plasmids (*ProOsNAS2:LUC* and *ProOsYSL15:LUC*) were transformed into agrobacteria with the pSOUP vector. The positive clones were selected on LB media with 25 μg/mL kanamycin and 10 μg/mL tetracycline. Agrobacteria were resuspended in infiltration buffer (10 mM MgCl_2_, 10 mM MES, pH 5.6) at a final OD600 of 1. Agrobacteria with the corresponding plasmids were mixed at a ratio (effecter1 : effector2 : reporter1 : reporter2=1 : 1 : 0.5 :0.5) and a final concentration of 0.2 mM acetosyringone was added. The agrobacteria were kept at room temperature for at least 2 h without shaking. Leaf infiltration was conducted in 3-week-old *N. benthamiana*.

The whole leaves infiltrated were used for ChIP assays. ChIP assays were conducted essentially according to previously described protocols (Saleh et al., 2008). ChIP assays were performed by anti-Myc antibody and anti-HA antibody respectively. To quantify OsFIT-DNA or OsIRO2-DNA binding ratio, qPCR was performed. The primers used for ChIP-qPCR are listed in Supplementary Table S2. For the quantification of each DNA fragment, three biological replicates were used. Each biological replicate contained three technical replicates.

### EMSA

EMSA was performed using a Chemiluminescent EMSA Kit (Beyotime). The recombinant GST-OsIRO2 and GST-OsFIT proteins were expressed in and purified from *E. coli*. Two complementary single-stranded DNA probes were synthesized and labeled by biotin at the 5’ terminus. Biotin-unlabeled fragments of the same sequences or mutated sequences were used as competitors, and the GST protein alone was used as the negative control. The sequences of the probes are shown in Supplemental Table S2.

### Immunoblotting

Protein samples were separated on a 12% SDS-PAGE and transferred to a nitrocellulose membrane. Target proteins on the membrane were detected using immunodetection and chemiluminescence. Signals on the membrane were recorded using a chemiluminescence detection machine (Tanon-5200). The antibodies used for western blot are as follows, mouse monoclonal anti-HA (Affinity Biosciences, Cat#T0050), mouse monoclonal anti-Myc (ABclonal, Cat#AE010), and goat anti-mouse IgG horseradish peroxidase (Affinity Biosciences, Cat#S0002).

### Agilent GeneChip Analysis

Four-day-old seedlings germinated in wet paper were transferred to solution culture with 0.1 mM Fe(III)-EDTA for five days. Then seedlings were transferred to solution culture without Fe or with 0.1 mM Fe(III)-EDTA for five days. Roots and shoots were separated and used for RNA extraction. GeneChip analysis was conducted by OE Biotech. Co. Ltd. (Shanghai).

## ACKNOWLEDGMENTS

We thank the Biogeochemical Laboratory and Central Laboratory (Xishuangbanna Tropical Botanical Garden) for assistance in the determination of metal contents. This work was supported by the Applied Basic Research Project of Yunnan Province (2017FB026) and the CAS 135 program (2017XTBG-T02).

## SUPPLEMENTAL DATA

**Supplemental Figure S1.** Self-activation of full-length OsIRO2 in yeast and similarity of AtFIT and OsFIT.

**Supplemental Figure S2.** Analysis of *ProOsFIT:GUS* lines.

**Supplemental Figure S3.** Generation and analysis of CRISPR/Cas9 edited mutants.

**Supplemental Figure S4**. OsFIT1 transcriptional regulation of Fe homeostasis genes.

**Supplemental Figure S5**. Analysis of *OsFIT* overexpression plants.

**Supplemental Figure S6**. Expression of *OsFIT* in the *iro2-1* mutant.

**Supplemental Figure S7.** Both OsIRO2 and OsFIT are required to regulate their targets.

**Supplemental Table S1.** Candidates interacting with OsIRO2 in yeast.

**Supplemental Table S2.** Primers used in this paper.

